# Dynamic clustering regulates activity of mechanosensitive membrane channels

**DOI:** 10.1101/553248

**Authors:** Alexandru Paraschiv, Smitha Hegde, Raman Ganti, Teuta Pilizota, Anđela Šarić

## Abstract

Experiments have suggested that bacterial mechanosensitive channels separate into 2D clusters, the role of which is unclear. By developing a coarse-grained computer model we find that clustering promotes the channel closure, which is highly dependent on the channel concentration and membrane stress. This behaviour yields a tightly regulated gating system, whereby at high tensions channels gate individually, and at lower tensions the channels spontaneously aggregate and inactivate. We implement this positive feedback into the model for cell volume regulation, and find that the channel clustering protects the cell against excessive loss of cytoplasmic content.

---

Both eukaryotic and prokaryotic cells harbour a phospholipid membrane packed with proteins, which enables separation of cellular content from the external environment. This physical barrier facilitates transport of signals and materials between the cell and its environment, thus sustaining life [1]. In addition, membranes of unicellular organisms separate the cell from the outside world, and need to be able to respond quickly and efficiently to sudden changes in the cell’s surroundings. One of the ways the membranes respond to external stimuli is by reorganising associated macromolecules [2]. A characteristic example of such a behaviour are membrane mechanosensitive channels (MSCs), which respond to mechanical cues from the cell’s surrounding, and are central to senses of hearing, balance, and touch, as well as for ensuring cell osmotic homeostasis [3–5].

The best studied MSCs are those of bacterium *Escherichia coli*, whose role is to protect the cell against sudden drops in the environmental solute concentration, so called hypoosmotic shock [6, 7]. Upon hypoosmotic shock water rushes into the cell, resulting in the cell swelling and increased tension in the bacterial envelope, which apart from the protein-filled phospholipid membranes, consists also of a stiffer material called the cell wall [6, 7]. If left unchecked, this pressure can lead to cell death by rupturing the envelope [8, 9]. To prevent it, a portfolio of MSC in *E. coli’s* inner membrane act as “pressure release valves” that open and create a nano-sized pore at the centre of the protein. This in turn enables solute and water efflux, reestablishing desired osmotic pressure inside the cell [10–12]. This response is fast and solely regulated by the membrane tension and chemical potential of water and solutes [12].

Bacterial MSCs consist of closely-packed trans-membrane helices connected by loops [13, 14]. Driven by membrane tension, the helices are thought to tilt with respect to one another, creating a space between them (up to 3 nm in diameter) for small solutes to non-selectively pass through [15]. Recent studies debate the existence and the role of spontaneous clustering of one of the MSCs found in *Escherichia coli*, MSC of large conductance (MscL) [16, 17]. Indeed, membrane clustering appears to be a common mechanism in cellular signaling, and has been observed for many transmembrane proteins and signaling receptors [18]. Clustering of MSC *in vitro* has been shown to result in collective, non-linear gating behavior [16], suggesting that it could tamper with cell’s passive response during a hypoosmotic shock recovery. However, assessing the extent of MscL aggregation via imaging techniques *in vivo* has proven to be difficult due to the potential artifacts of the MscL tags on the process [17], and hence opens a need to explore orthogonal ways to investigate the aggregation phenomenon. Here, by developing a minimal computer model of MSCs embedded in a fluid membrane, we investigate the physical mechanisms behind the MSC cluster formation and cooperative gating, and their implications on cell-volume regulation.

Guided by the known structures of single isolated MSCs [13, 14, 19], we built a generic MSC model out of rod-shaped subunits connected by weak springs (Fig. 1a). While bacterial MSCs possess varying number of repetitive helical sub-units [7], without loss of generality, we choose to include five rod-shaped subunits. Each rod is made of seven core hydrophobic, and two hydrophilic head beads (Fig. 1a). The rods are longer than the membrane thickness to reproduce a positive hydrophobic mismatch of ~ 0.5nm between the protein and the lipid layer found in structural studies [13]. The lipid bilayer is described by a previously published three-beads-per-lipid model [20] (Fig. S1 and S2). The single rod diameter is twice the radius of a lipid bead, and the inner part of the channel is lined with hydrophilic beads to prevent lipids from overflowing inside the channel. Finally, to be able to include direct inter-protein attractions, an attractive patch of beads is added on the external side of each rod. Hypoosmotic shock is generated by placing a gas of inert volume-excluded “solute” beads on one side of the membrane. The collisions of the solute beads with the membrane create membrane tension, which is linearly proportional to the solute concentration difference across the membrane (Fig. S5). For further details on simulations see Supplementary Information.

**FIG. 1:**
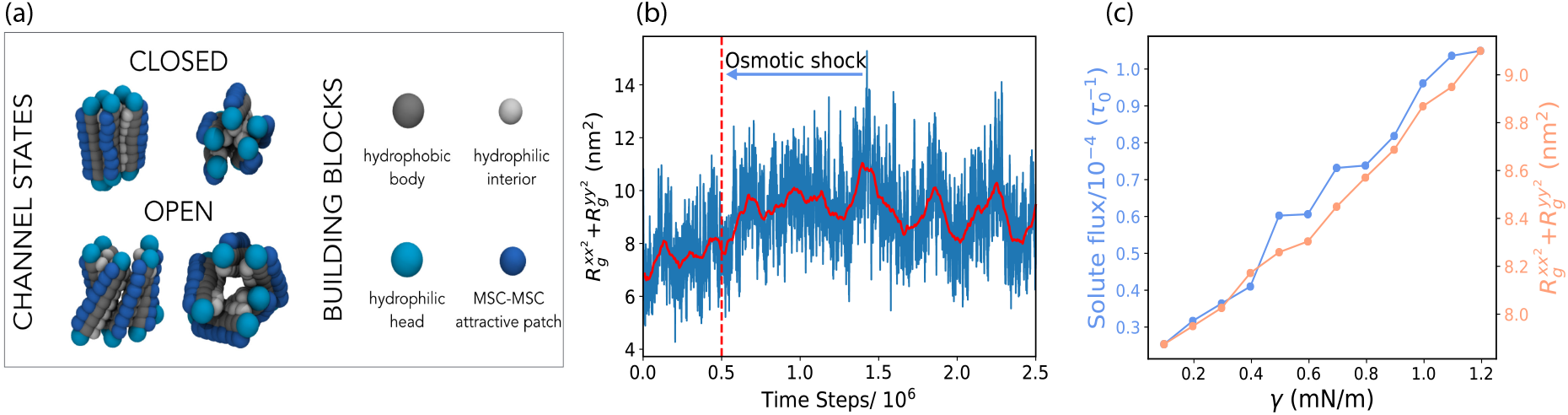
Coarse-grained model and single channel properties. (a) MSC is presented as a collection of rods connected by weak springs. Each rod consists of overlapping hydrophobic beads (depicted in gray) and hydrophilic heads (in cyan), and is ~ 10 nm long. Channel inside is lined with hydrophilic stripes (silver). Explicit inter-channel attractions can be turned-on via an external hydrophobic patch (dark blue). (b) Pore size oscillations of a single MSC. The dashed vertical red line marks the occurrence of an instantaneous osmotic shock corresponding to the membrane tension of *γ* = 1.2 mN/m, which leads to an increase in the average pore size. The solid red line represents the moving time average (window size 10^5^ time steps). (c) The variation of the pore size, quantified by the in-plane components of the MSC radius of gyration tensor 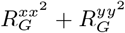, and subsequent solute flux though the channel as a function of the membrane tension.

We first focus on the behaviour of a single MSC. Application of a hypoosmotic shock causes an increase in the membrane tension and area. Since the transmembrane components of the MSC interact attractively with the hydrophobic layer of the membrane, they maintain contact with the expanding lipid bilayer. Consequently, channel rods tilt with respect to one another, resulting in the overall lateral expansion of the channel, as shown in Fig. 1 and Video 1. To quantify the channel pore size we measure the in-plane components of the MSC radius of gyration tensor (see Supplementary Information). We find that the pore size oscillates stochastically, and that the application of hypoosmotic shock leads to an immediate increase in the pore size (Fig. 1b), allowing for the passage of the solutes and channel gating (Video 2). As shown in Fig. 1c, the pore size and the flux of solute through the pore increase with the increase in the shock magnitude. For the purpose of our analysis solely, we chose *γ* = 0.45 mN/m as the threshold tension for the pore opening; channels whose pore size is above 8.2 nm^2^ we consider as open, while those below this size we deem as closed (Eq. (S5)).

We now analyze the behavior at multiple MSCs interacting only via volume exclusion and effective membrane-mediated interactions. Fig. 2 shows the gating properties of such MSCs as a function of the number of channels in the system. In this case we did not observe any channel clustering and it is evident that each channel behaves independently. Indeed, we find that the membrane-mediated interactions between fluctuating channels in our system are negligible (Fig. S7). Rigid symmetric inclusions of the same hydrophobic mismatch in our model experience attraction of ~ 0.5kT (Fig. S8), in agreement with previous simulation studies [21–23]. redAs we did not observed any MSC clustering due to pure membrane-mediated interactions, we chose a top-down strategy. We know that: (i) MSC aggregation has been reported *in vitro* [16], and *in vivo* while working with MSCs labelled with a small covalent dye [17], (ii) direct inter-protein interactions, such as polar and electrostatic interactions and packing of small apolar side chains [24–29], can lead to attractions of trans-membrane proteins [18]. In addition, local lipid phase separation around the protein can yield effective inter-protein attraction and protein aggregation. Therefore, to drive MSC aggregation we included weak direct inter-protein attractions, which we modelled via an attractive stripe on the outer side of the channel (Fig. 1a).

**FIG. 2:**
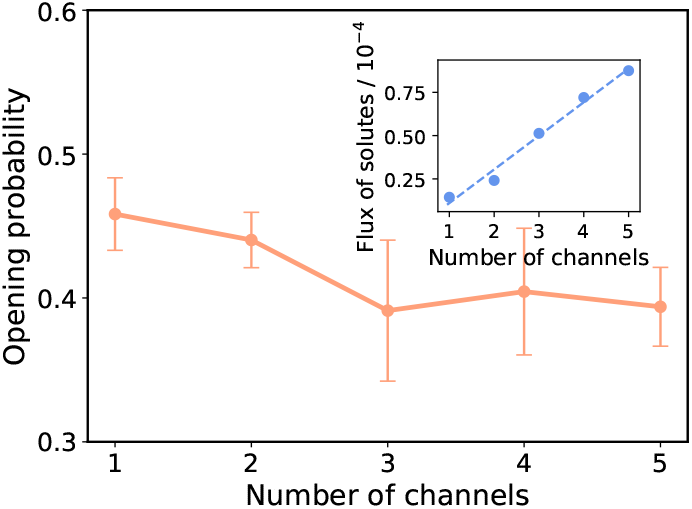
Multiple channels interacting via membrane-mediated interactions only do not cluster and hence gate independently. The probability of channel pore opening versus the number of channels present in the system. Inset: total flux of solutes through the channels versus the number of channels in the system. Channel area fraction ranges from 0.03 (*N* = 1) to 0.15 (*N* = 5).

Incorporating weak direct inter-protein attraction leads to the assembly of MSCs into small clusters of sizes between 2 and ~ 15 MSCs, and we now find that the clusters exhibit strong cooperative gating. Fig. 3(a) shows that the pore sizes of two attractive channels varies as a function of the separation between them. Sharp decrease in the pore sizes at ~9.5 nm of inter-channel separation corresponds to the cooperative closure of individual MSC (Video 3). The reason for this is purely geometrical: two closed channels can achieve larger contact area between them, maximizing their attraction. For multiple channels diffusing in the bilayer we observe dynamic rearrangement and aggregation into larger clusters that leads to decreased gating activity per channel, which scales with the cluster size (Fig. 3b). The clusters are dynamic in nature, whereby individual channels within the clusters oscillate between the closed and open states, can move within, leave the cluster, or join another. Since the channel activity depends on the number of neighbours, individual channel activity within a single cluster is consequentially inhomogeneous. The channels on the cluster interior will on average gate less than the channels sitting at the aggregate rim (inset in Fig. 3b). The average channel activity will hence depend not only on the cluster size, but also on its shape.

**FIG. 3:**
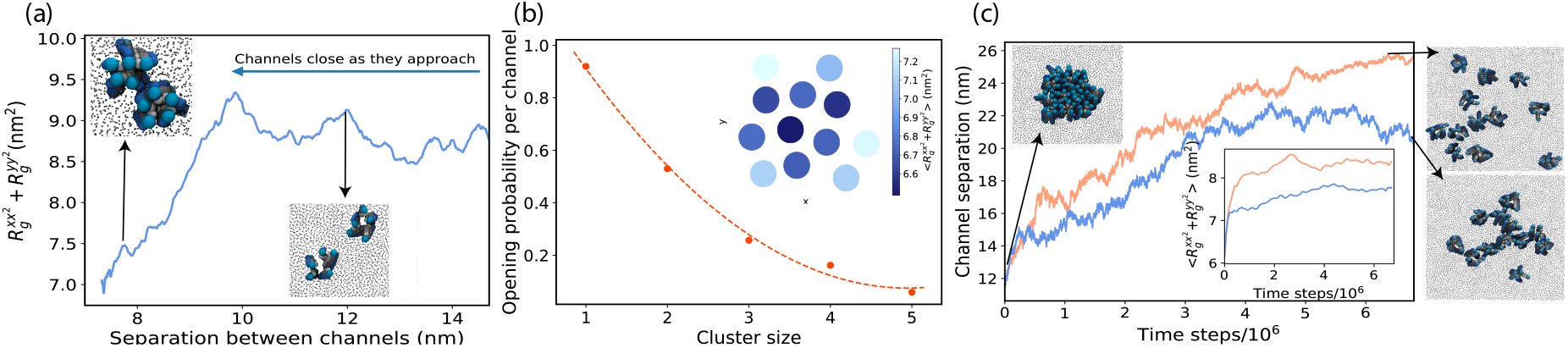
Channels interacting via explicit attractive interactions exhibit cooperative gating. (a) Average pore size of two MSCs as a function of distance between them (*γ* = 1.30 mN/m). (b) Opening probability per channel versus aggregate size (*γ* = 1.70 mN/m and the channel area fraction is 0.16). Inset: The average pore size of channels within a single cluster formed of twelve MSCs. (c) The average distance between the channels increases in time and is larger for higher membrane tension (*γ* = 1.70 mN/m, shown in orange versus *γ* = 0.70 mN/m, shown in blue) Far right: The snapshot of the cluster configuration in the last time-frame. Inset: The probability of channel opening increases as the simulation progresses. In all the subfigures *ε*_*protein*−*protein*_ = 0.9kT.

We now perform a computational experiment to mimic the situation in which a bacterial cell, living under quiescent conditions, encounters a sudden hypoosmotic shock. We start our simulation with a membrane that contains channels all aggregated into a single cluster. We then applied a sudden tension of 1.5 mN/m, and monitored the system in time. As a control, the same simulation was repeated at zero tension (Fig. 3c). We find that the high magnitude shock breaks up the MSC cluster into individual channels, switching the system from the clustered to the mixed state (Video 4 and 5). Such isolated channels open with higher probability (inset in Fig. 3c), enabling efficient gating at high membrane tensions, when it is needed most. On the contrary, at low tensions the channels remain in a cluster, albeit the cluster shape dynamically elongates (Fig. 3c).

These findings suggest that the spontaneous formation of liquid-like MSC clusters enables an additional level of control over their gating and signal transduction. This control is implemented in the system in a passive way, hard-wired in the system’s physical properties. On average, single channels are more closed at low membrane tensions, making the channels more aggregation-prone, which in turn further deactivates their gating. When the cell encounters a hypoosmotic shock, the membrane tension increases and channels open, making them less aggregation prone, which results in spontaneous dispersion of clusters and further opening of individual channels. The positive feedback between the membrane tension and the cluster formation hence dynamically adjusts the extent of channel clustering, as well as their gating properties.

We now include the observed effect of channel clustering into our previously developed continuum model of *E. coli* cell volume recovery upon hypoosmotic shock [12]. Experimentally, we observed total cell volume expansion within seconds after the hypoosmotic shock, followed by a period of slower, minutes-long volume recovery (despite the fact that MSCs open on miliseconds time scales) that exhibits a characteristic “overshoot” below the value of initial volume (Fig. S11). Our continuum model explained the slow recovery and the volume recovery overshoot by considering the change of the cellular volume (*V*_*n*_) and solute concentration in time. The volume changes, and consequentially the cell membrane tension, are governed by the flux of water, proportional to the difference between osmotic pressure and Laplace pressure on the cell wall. Solute concentration changes are governed by the diffusive fluxes through the MSCs, enhanced by the tension build up (see Supplementary Information).

Thus, when MSCs are open the solute flux through the cell membrane increases, which was described in the model with a single fitting parameter that characterizes channels as either open or closed (Eq. (S10)). To link our coarse-grained model predictions with the continuum model we now replace that parameter with a continuous function, capturing that the channel clustering: *(i)* decreases at higher membrane tensions, *(ii)* decreases opening probability per channel and *(iii)* increases for higher MSCs surface fractions. The introduced function hence depends on the number of channels in the cluster (*N*), bilayer tension (*γ*), and the channel surface density (*ρ*), and any constants are fixed by fitting to the results of the coarse-grained model (see Supplementary Information).

Next, we fit the experimental volume trace to the master equations (Eq. (S23) and (S24)) that describe the changes in cellular volume and solute concentration with the results of the coarse-grained model incorporated (Fig. S11). The fit enables us to predict the dynamics of cluster aggregation and disaggregation as the cell volume expands and recovers after hypoosmotic shock, Fig. 4a. Probability of observing channels as monomers (*N* = 1) and as aggregates (shown for *N* = 5) is given as a color scale for each time point post-hypoosmotic shock, showing that larger clusters are less likely to form at the point of maximum volume expansion (largest tension), and more likely to form as the volume recovers and the membrane tension decreases. This gives a clear prediction of our model, which can be tested by imaging the extent of the channel clustering in the membrane at different times post-hypoosmotic shock.

**FIG. 4:**
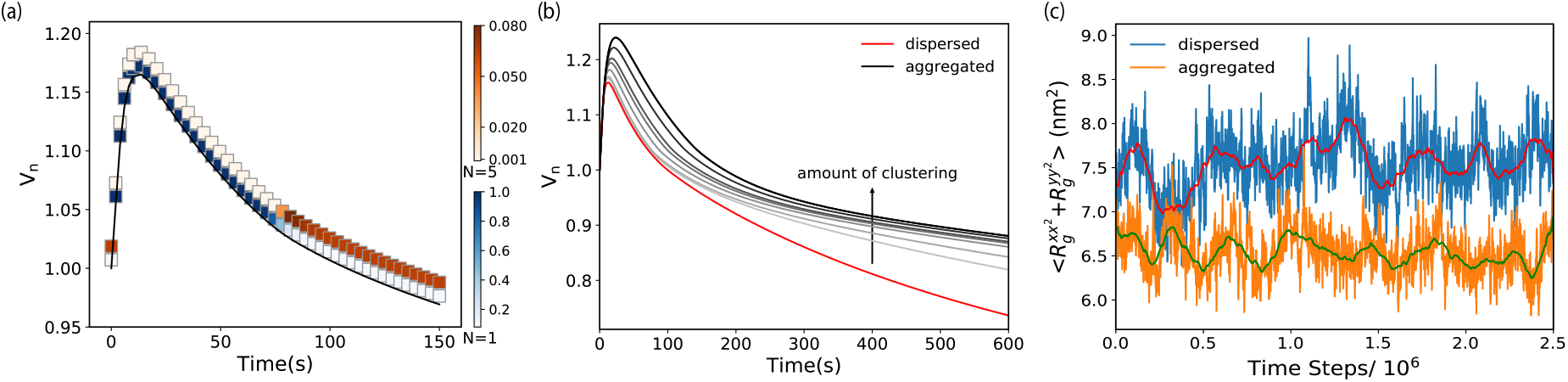
Clustering regulates channel closing to overcome leaky cell membrane. (a) Dynamics of channel clustering during cell volume recovery (gray faded line). Normalized cell volume, *V*_*n*_, upon 0.96 Osmol hypoosmotic shock and 0.5% channel packing fraction. Colorbars: Probability of finding a single isolated channel (blue) and a cluster of 5 channels (orange).(b) Effect of increasing the extent of clustering (here by changing protein-protein attractions), on cell volume dynamics (grey and black lines) in comparison to dispersed channels (red line) for the same conditions as in (a). (c) Oscillations in the average MSC pore size at zero membrane tension for a system of 12 MSCs in a dispersed (blue line) and aggregated (orange line) states. Red and green lines show the moving time average in each case (10^5^ time steps window).

To demonstrate the consequence of MSCs clustering on the cell volume recovery, we show what the volume dynamics would look like if the channel were prevented from clustering and if the clustering differed from that of the fitted data (Fig. 4b and Fig. S12). This allows us to see that, within a specific range, channel clustering can reduce the volume “overshoot” commonly found upon recovery, without jeopardizing channel opening at the point of maximum tension. Fig. 4b and Fig. S13 show that further decreasing the overshoot by MSC clustering can lead to detrimental increase in the maximum tension in the cell envelope, suggesting that the channel clustering is finely-tuned in the cell. Our prediction on the role of clustering for the cell volume regulation can be probed experimentally by tracing the volume recovery post-hypoosmotic shock for different extents of the clustering. The MSC clustering can be enhanced by tagging the channels with fluorescent proteins [17] or modulated by expressing the channels to different levels.

In conclusion, we showed that spontaneous aggregation of mechanosensitive membrane channels results in liquid-like clusters that exhibit lower gating activity than dispersed clusters. Our findings align well with the study by Grage et al. [16] in which the aggregation of *E. coli* MscLs reconstituted in lipid vesicles led to a significant decrease in the total gating activity. The patch-clamp experiments in [16] showed that a number of active channels in a patch was consistently lower than the total number of channels. These results were further reinforced by small angle neutron scattering measurements of the total membrane area increase when the channels were open, which was smaller than what would have been expected if the channels were behaving independently.

Previous continuum models predicted that, due to hydrophobic mismatch, membrane-mediated interactions between perfectly rigid symmetric mechanosensitive channels will lead to their collective opening [30–34]. This is not what we have observed in our coarse-grained model. Our channels are not perfectly symmetric or rigid, but can dynamically acquire different conformations, which can render membrane-mediated interactions between two channels both attractive and repulsive. It is likely that this effect erases any membrane-mediated interactions. Interactions between perfectly rigid inclusions of hydrophobic mismatch of 0.5 nm in our model are attractive, albeit weak (Fig. S8). It is possible that the high stiffness of the coarse-grained lipids prevents lipid stretching needed for membrane-mediated interactions. Nevertheless, since membrane-mediated interactions due to hydrophobic mismatch have never been directly experimentally quantified, it is hard to assess their importance in driving MSC aggregation observed in experiments [16, 17]. There is however a growing body of evidence that direct protein-protein interactions drive aggregation of transmembrane helices, and also stabilize helix-helix interactions within a single protein [24–29]. It is likely that the same forces could also drive weak helix-helix interactions between different proteins should they be found close to each other.

We demonstrated that coupling between the membrane tension, channels’ conformational change and clustering produces a controlled gating system, whose positive feedback is encoded purely in the system’s physical properties. Based on these results, we predict the effects of the feedback on the cell volume regulation. We suggest that MSC aggregation serves to protect the cell from excessive gating, both in steady-state and during its post-shock volume recovery. Indeed, our simulations show that isolated channels have a non-zero probability of gating even at zero tension (Fig. 4c). This agrees with experimental characterisation of a single-channel gating, where it is evident that the channel opening does not follow a sharp step function [35]. Hence, if MSCs are over-expressed, e.g. when bacteria grow under hyperosmotic conditions or when they enter stationary phase [36], the probability of single channel gating even under quiescent conditions would be sufficiently high tovsignificantly increase the effective membrane permeability to ions, making it hard to maintain electrochemical gradients across the cell membrane, which serve as one of the main energy sources for the cell [37–42]. Furthermore, loss of volume by 8-10% has been experimentally reported to lead to the loss of turgor pressure that the cell actively maintains [43]. Thus channel aggregation, which is more pronounced at higher channel numbers, could be a natural self-defence mechanism of bacteria against unnecessary gating, contributing to bacterial survival, especially in scares environments (alike post-hypoosmotic shock). The MSC model developed identifies the basic physical mechanisms behind mechanosensing of membrane channels. Due to their generality, our results can also be helpful in guiding the design of synthetic nanomechanosening systems [44] and artificial membrane channels [45, 46].

## Acknowledgments

We thank Samantha Miller, Bert Poolman, and the members of Šarić and Pilizota laboratories for useful discussion. We acknowledge support from the Engineering and Physical Sciences Research Council (A.P. and A.Š.), the UCL Institute for the Physics of Living Systems (A.P. and A.Š.), Darwin Trust of University of Edinburgh (H.S.), Industrial Biotechnology Innovation Centre (H.S. and T.P.), BBSRC Council Crossing Biological Membrane Network (H.S. and T.P.), BBSRC/EPSRC/MRC Synthetic Biology Research Centre (T.P.), and the Royal Society (A.Š.).

