## Supplementary Information for "Dynamic clustering regulates activity of mechanosensitive membrane channels"

### Supplementary Information: Dynamic clustering regulates activity of mechanosensitive membrane channels

This Supplementary Information provides details related to implementation of the coarse-grained model for lipids and mechanosensitive channels (Section I), details regarding the simulation set-up (Section II), details on the model of cell volume dynamics during hypoosmotic shocks (Section III) and supportive results and figures in Section IV.

#### I. COARSE-GRAINED MODEL

##### A. Lipids

The lipid bilayer was simulated using a highly coarse-grained model developed by Cooke and Deserno [1]. In this model, each lipid is made of three beads: one hydrophilic "head" bead and two hydrophobic "tail" beads. The beads are joined together by finite extensible nonlinear elastic (FENE) bonds described by the bond potential $V_{bond}(r) = -\frac{1}{2}k_{bond}r_{\infty}^2 \log\left[1-\left(\frac{r}{r_{\infty}}\right)^2\right]$, where $r$ is the distance between atoms and $r_{\infty}$ is the maximum bond extent, in this case taken to be $1.5\sigma$. We used $k_{bond} = 30kT/\sigma^2$. $\sigma$ is the Lennard-Jones unit of length and corresponds to approximately 1 nm. The angle between the three beads is enforced by a harmonic angular potential $V_{angle}(r) = k_{angle}\left(\theta - \theta_0\right)^2$, with $k_{angle} = 5kT/deg^2$ and $\theta_0 = \pi$.

The intermolecular interactions between the lipid beads are determined by a perturbed Weeks-Chandler-Andersen (WCA) potential [2]. All beads interact via a truncated-shifted Lennard-Jones potential described in equation (S1). The parameter b gives the relative size of the lipid beads and $r_c$ is a cutoff parameter. In our simulations we used $\varepsilon = 1kT$, $r_c = 2^{1/6}\sigma$, $b_{head,head} = b_{head,tail} = \sigma$, $b_{tail,tail} = 0.95\sigma$ as described in [1].

$$V_{rep}(r,b) = \begin{cases} 4\varepsilon\left[\left(\frac{b}{r}\right)^{12} - \left(\frac{b}{r}\right)^{6} + \frac{1}{4}\right], & r \le r_c \\ 0 & r > r_c \end{cases} \tag{S1}$$

An additional attractive potential is enforced between the hydrophobic tail beads [1]. The membrane is kept intact only through the attractive interactions between the hydrophobic tail beads. This perturbative potential is described in equation (S2):

$$V_{attr}(r) = \begin{cases} -\varepsilon, & r < r_c \\ -\varepsilon cos^2\frac{\pi(r-r_c)}{2w_c}, & r_c \le r \le r_c + w_c \\ 0, & r > r_c + w_c \end{cases} \tag{S2}$$

The cosine term is meant to soften the potential which makes the membrane more fluid and ensures the self-assembly and membrane's structural integrity. The adjustable parameter $w_c$ controls decay range of the potential. In our simulations, we used $w_c = 1.4\sigma$. All the parameters were chosen such that the membrane is positioned well into the fluid phase of the phase diagram reported in the original paper [1].

##### B. Mechanosensitive Channels

We designed a similar coarse-grained model for the mechanosensitive channel of large conductance (MscLs). A single channel is made from five rod-shaped subunits, inspired from the protein's oligomeric structure (Fig. S1). Each subunit mimics the alpha-helices from the transmembrane domains TM1 and

TM2. A representation of a channel is given in Figure S1. The rods are interconnected by a series of weak harmonic bonds of spring constant $k_{spring} = 1.5kT/\sigma^2$, with an equilibrium length set to $r_{eq} = 2.5\sigma$. Each rod is made of seven core hydrophobic beads and two hydrophilic head beads. The rods are slightly longer than the membrane thickness in order to replicate the hydrophobic mismatch between the protein and the lipid bilayer. All the beads are connected by a strong harmonic potential ($k = 100kT/\sigma^2$, $r_{eq} = 2.5\sigma$) meant to ensure the channel's rigidity. The channel is kept straight by a similar angular harmonic potential $k_\theta = 100kT/deg^2$, $\theta_0 = \pi$.

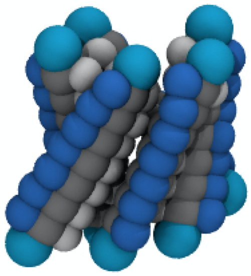

FIG. S1: **Coarse-grained model of a single mechanosensitive channel**. There are four different types of beads in a channel: hydrophilic head beads (cyan), hydrophobic core beads (grey), hydrophilic interior beads (silver), attractive MscL-MscL beads (blue).

The channel head and core beads interact with the lipids and with themselves through the same potential described in equations (S1) and (S2). The size of these protein beads was chosen to be $2\sigma$, giving the relative parameters $b_{protein,protein} = 2\sigma$ and $b_{protein,head} = b_{protein,tail} = 1.5\sigma$. The head beads interact only with WCA potential (S1). The interactions of the core beads contain both the attractive and repulsive parts of the potential (S1 and S2). These beads are hydrophobic so they are attractive towards the lipids and keep the channel embedded within the membrane. Attempts of using solely a Lennard-Jones potential for the channel-membrane interaction, without the soft cosine tail, led to unstable structures that quickly fractured at the point of contact between the protein and the lipids. This suggests that a careful tuning in the interaction between proteins and membrane is crucial for functionality of MscLs.

A hydrophilic patch was added on the channels' interior side to prevent lipids from overflowing inside (silver beads in Figure S1). They interact with the lipids purely through the repulsive part of the Lennard-Jones potential. Their diameter was chosen to be $0.5\sigma$ and the potential depth was taken to be $\varepsilon = 50kT$. Since these beads are not attractive to the lipids, they prevent them from flowing inside the channel. Another attractive patch of beads is added on the external side of the channels (blue beads in Figure S1). This patch drives the interaction between proteins via a long-range Lennard-Jones potential (with a cutoff of $3\sigma$) with adjustable depth $\varepsilon_{protein,protein}$. Both these types of beads were connected with the main core beads via a spring with spring constant $k = 1.5kT/\sigma^2$ and equilibrium length $r_{eq} = 1.5\sigma$. The attractive patch is essential to drive the aggregation of channels into clusters and controls the cooperative gating effects described in the paper.

#### II. SIMULATION DETAILS

The channels were inserted vertically in a lipid membrane placed in the centre of the simulation box. The lipids were initially organised into a square patch on a lattice which further reorganises in a typical bilayer-like structure. The membrane was tethered to the centre of the simulation box with a weak spring of spring constant $k = 1.0kT/\sigma^2$ to prevent its drift under the collisions with the solutes. The simulation box is cuboidal in shape with the initial dimensions $L_x = L_y = \frac{1}{3}L_z$. The side length of the box is allowed to fluctuate in the $x$ and $y$ directions with periodic boundary conditions applied. The simulation box was kept fixed in size in the $z$ direction with weak Lennard-Jones walls at both ends, and with the wall depth $\varepsilon = 1.0kT$ .

Our system contained 6024 lipids, each of which is made of 3 particles. We simulated at most 12 channels, where each is made of 115 particles. Number of solute particles was varied between 0 and 1200, where each solute is made of 1 particles. Hence the largest simulation we ran contained 20,652 particles.

The simulations were run in the isoenthalpic-isobaric ($NPH$) ensemble with zero lateral pressure $P_x = P_y = 0$. The membrane tension is controlled by inserting external osmolyte particles that collide with the membrane. This mechanical interaction between the solutes and the membrane leads to a subsequent expansion of the membrane in the $xy$ plane and a small thinning in the $z$ direction. In order to replicate the stochastic dynamics of the real system, we used a Langevin thermostat with the friction coefficient $\eta$ set to unity: $\eta = m/\tau_0$, where $m$ is the particle mass (set to unity for all particles) and $\tau_0$ the simulation unit of time. The simulation time step was taken to be $\tau = 0.008\tau_0$, where $\tau_0$ is the simulation unit of time. We used the LAMMPS molecular dynamics package to run the simulations [3] and the VMD package to visualise the trajectory files [4].

##### A. Membrane tension

Membrane tension is controlled by the number of solute particles impinging on the membrane. The aim of these particles is to mimic the chemical concentration gradient that a channel is exposed to during an osmotic downshock. The solute particles are modelled as hard spheres of diameter $0.5\sigma$ that interact with the other particles in the system (including themselves) only through the WCA potential with $\epsilon = 50kT$ (SI). The size of the solutes was chosen carefully so they can flow through the open channels, but not through channels with more contracted conformations or through the membrane itself.

The surface tension was mapped to the number of solute particles as follows: A number of solute particles, $N_{solutes}$, were placed in a box containing a square patch of a lipid membrane of the side length $40\sigma$ (in the absence of any mechanosensitive channels). All the solute particles were placed on the same side of the simulation box with respect to the membrane patch. The membrane tension, $\gamma$, was calculated by integrating the normal and tangential components of the pressure tensor using the relationship S3 as described in [5]:

$$\gamma = \int [P_{zz} - \frac{1}{2}(P_{xx} + P_{yy})]\mathrm{d}z \tag{S3}$$

Further results are given in Section IV.

##### B. Channel radial expansion

The radial expansion of a channel is tracked using the $xx$ and $yy$ components of the gyration radius tensor as described in equation

$$R_G^{xx^2} + R_G^{yy^2} = \frac{1}{M}\sum_i m_i(r_{ix} - r_{CMx})^2 + \frac{1}{M}\sum_i m_i(r_{iy} - r_{CMy})^2 \tag{S4}$$

where $i$ is the index of the given protein bead, $m_i$ is its mass and $M$ is the total channel mass. $r_{ix}$ and $r_{iy}$ are the Cartesian coordinates of the particles in the $x$ and $y$ directions. $r_{CMx}$ and $r_{CMy}$ are the coordinates of the channel's centre of mass.

We assigned a state function to the channel, depending on the corresponding gyration radius components. The channel can be considered either "open" or "closed" depending on its radial expansion. As such, the state of a channel is defined as:

$$\text{Channel state} = \begin{cases} \text{open}, & \text{if } R_G^{xx^2} + R_G^{yy^2} \geq R_t \\ \text{closed}, & \text{otherwise} \end{cases} \tag{S5}$$

where $R_t$ is the a threshold value above which the channel is considered open. In this case, it was chosen to be 8.2 $nm^2$, relying on the sharp transition observed in the pore size with increasing membrane tension (Fig. 1c in the main text). The opening probability was further calculated as $P_{\text{open}} = \text{open states/total states}$, where the state of the channel is recorded at every time step.

##### C. Single channel

A single channel was inserted in a square membrane patch of side length $40\sigma$. The system was equilibrated for 500,000 time steps and a hypoosmotic shock was applied by inserting $N_{solutes}$ solute particles in the system. Multiple simulations were run by varying $N_{solutes}$ between 0 and 1000 in increments of 100 particles. The simulations were run for 2.5 million time steps each. The radial expansion of the channels was recorded at every step during the simulation. Figure 2(a) shows how the radial expansion of the channel is affected during a downshock corresponding to a tension of approximately 1 $mN/m$. The fluctuations of the radial expansion are increased to higher values immediately after the application of the downshock, indicating that the channel acts like a pressure sensor capable of sensing small changes in the surrounding membrane tension.

##### D. Non-interacting channels

$N$ channels were inserted inside a membrane patch, where $N = 1, 2, 3, 4, 5$. $N_{solutes} = 800$ were added to the system. Both the probability of opening and the flux of solutes (the flow rate of the solute through the membrane) were recorded for 2.5 million time steps, similarly to the single channel simulation. Five different random seeds in the initial channel geometry were used for each simulation. Results are presented in Fig. 2 of the main text. It was revealed that the number of channels had no influence on the opening probability of individual channels and that the flux of solutes is linear in the number of channels. These results indicate that the channels do not influence each other's gating properties in the absence of a direct interaction. The model does not adequately capture any potential membrane-mediated interactions. Therefore, an attractive patch (blue beads, Fig S1) was added in order to investigate the effect of nonspecific protein interaction on gating.

##### E. Interaction between two channels

To measure the influence of two adjacent channels on one another, we fixed them in the membrane patch at a $d = 10\sigma$ distance with very weak springs of elastic constant $k_{\text{spring}} = 0.5kT/\sigma^2$. The weak springs serve to maintain the channels in each other's proximity and also limit their lateral diffusion. The interaction between the attractive patches located on the two channels was increased from $\varepsilon = 0kT$ to $\varepsilon = 2.0kT$ in increments of $0.1kT$. The influence of two channels interacting on their radial expansion is described in Fig. 3a of the main text. There is a clear tendency for adjacent interacting channels to reduce each other's radial expansion, and force each other to adopt a more closed conformation. Conformational changes require a higher activation energy necessary to overcome the attractive homophilic protein interactions. The effect of interaction energy on the opening probability of adjacent channels is described in section IV.C. The simulations with two channels were run for 2 million steps and five random seeds were used for the initial geometry. The effect of protein interactions was studied at two different membrane tensions 1.30 mN/m, 0.70 mN/m.

##### F. Aggregates of MscLs

Twelve MscLs were inserted in a membrane patch of side length $60\sigma$ at random initial locations. The membrane patch was equilibrated for 500,000 time steps. Two different membrane tensions (1.70 mN/m, 0.70 mN/m.) were applied. The depth of the interaction potential between the attractive protein patches was set to be $\varepsilon = 1.0kT$ to drive the aggregation of proteins into clusters (Fig. S2). A clustering algorithm was used to keep track of the MscL clusters formed during the simulation as follows: A channel was considered as being part of a cluster if it was found at a distance less than $10\sigma$ from a neighbouring channel, the distance at which two channels start interacting. A single channel radius is $\approx 4\sigma$. Changing the threshold to $9\sigma$ and $12\sigma$ did not make a significant difference in the distribution of the cluster sizes. The simulation was run for seven million time steps. The pore size was tracked at every time step and the average opening probability per channel per cluster was computed as a function of the cluster size (Fig. S3).

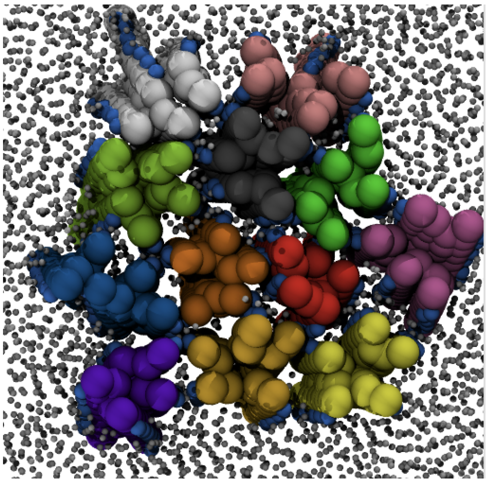

FIG. S2: **A cluster with twelve MscLs**, each presented with a different colour. The attraction between channels leads to the eventual aggregation of channels into large ensembles which show different gating behaviour from the individual channels. The attractive patches on the channels are coloured in light blue and the lipids in grey. Some of the channel beads are shown in a larger size than they are.

The formation of dimers was tracked throughout the simulation and the average interaction energy between the channels within a dimer was calculated at every time step. This allowed the estimation of the tension-dependent interaction between two channels. Further details are given in section IV.C.
For simulations that started with pre-formed clusters (Fig. S2), a single cluster was formed by slowly pulling all the twelve channels toward the centre of the membrane with weak springs. The springs were then released, but the attractive interaction between channels kept the cluster stable as long as no tension was applied to the membrane (Fig. 3b inset). A large osmotic downshock was then applied (corresponding to approximately 1.70 mN/m in one simulation and 0.70 mN/m in another). Both the separation between the channels and the pore size were tracked during seven million steps (Fig. 3c).

#### III. CELL VOLUME DYNAMICS DURING HYPOOSMOTIC SHOCK

##### A. A brief description of the previous model

We implement our previously developed model to deduce cellular volume ($V$) and solute ($n_i$) dynamics of *Escherichia coli* [6] upon a hypoosmotic shock. Water flux ($j$) across the cell membrane depends on the difference between the osmotic pressure ($\Pi$) and the Laplace pressure ($P$) of the cell wall, $j \sim -\pi - P$. Where $\pi$ is defined by Morse Equation given as below:

$$\Pi = -\phi(c_i - c_e)RT \tag{S6}$$

The molar osmotic co-efficient ($\phi$) is set to 1 because we osmolarities of solutions measured with an osmometer. Constants $c_i$, $c_e$, R and T are cytoplasmic and media solute concentrations, ideal gas constant and thermodynamic temperature, respectively. To derive $P$ we consider cell wall elasticity that has been experimentally demonstrated to exhibit stress-stiffening, thus $E = E_0(\dot{P}/P_0)^\kappa$ [7], where $E_0$ and $P_0$ are the pre-shock steady-state elasticity and pressure and $\kappa$ is the set to 1 (experimentall it was measured as 1.22) [6, 7]. We then write:

$$E = \frac{True \quad stress}{True \quad strain} = \frac{d\gamma/l}{dr/r} = E_0 \frac{P}{P_0} \tag{S7}$$

where $\gamma$ is the membrane tension, and $r$ and $l$ are the radius of the cell and the thickness of the cell wall (note that we effectively merge the contribution from the wall and the membranes into one, thus $l$ can be considered the whole envelope thickness). Considering *E.coli* as thin cylinder, we can write $\gamma = Pr$. At pre-shock steady-state, osmotic pressure gives rise to Laplace pressure, $P_0 = RT\Delta c_0$, where $\Delta c_0$=0.04 Osmol/l is the pre-shock osmotically active solute gradient across the cell membrane determined based on experimental estimates of turgor pressure [6]. Solving equation S7 for $P$ gives:

$$P = e^{\sqrt[3]{\frac{10}{3}\pi}\frac{E_0 l (V^{\frac{1}{3}} - V_0^{\frac{1}{3}})}{\Delta c_0 RT V^{\frac{1}{3}} V_0^{\frac{1}{3}}}} \cdot \Delta c_0 RT \frac{V_0^{\frac{1}{3}}}{V^{\frac{1}{3}}} \tag{S8}$$

$V_0$ is the pre-shock cell volume which is experimentally found to be 1.338 $\pm$ $\mu m^3$ [6]. The molar water flux during an osmotic shock leads to change in cell volume and hence:

$$\frac{dV}{dt} = V_m j A_c = V_m K \cdot (-\Pi - P) \tag{S9}$$

$A_c$ is the surface area of the cell (considering the cell as a spherocylinder of 2:1 length to diameter ration with d=1$\mu$m). $V_m$ is the molar volume of water and $K$ is the effective water conductivity. Gating of MSCs upon a hypoosmotic shock increases the permeability of the cell membrane to water, which we capture with a constant $A$

$$\frac{dV}{dt} = (A+1) \cdot V_m K \cdot (-\Pi - P) \tag{S10}$$

$A = 2$ means 2 times higher conductivity compared to the cell membrane with closed MSCs (when $A = 0$). The channels are set open if $V$ is greater than the volume threshold of opening ($V_{th}$). To deduce the dynamics of cytoplasmic solute concentration we allow the diffusive flow of solutes through the open MSCs, as well as take into account the build up of Laplace pressure in the cell wall due to the large influx of water:

$$\frac{dn_i}{dt} = -A \cdot \left( V_M K \cdot \frac{n_i}{V} \cdot P + D_s N_{MSC} \cdot a_{MSC} \cdot \frac{\frac{n_i}{V} - c_e}{l_M V_0} \right) \tag{S11}$$

where $N_{MSC}$ is the sum of all the MSCs in the cell, $a_{MSCL}$ is the total MSCs pore area, $D_s$ is the average diffusion constant of osmotically active solutes and $l_m$ is the thickness of cell membrane, both have been experimentally estimated and we use the same values as previously [6]

.

##### B. Including MscL clustering into the model

The channel we built for our course-grained simulations (Fig. S1) is based on the structure of MscL. In a wild type *E. coli* strain there are 7 different MSCs [8]. However, for simplicity we will apply the results of the course-grained simulations to them all and will for now exclude any differences. We thus use MSCs and MscLs interchangeably from here on. The gating activity of the channels will depend on the probability of opening of single channels and the extent of their clustering, which both depend on the membrane tension and the number of channels in the system. To include the effect of channel clustering on cell volume, we introduce a parameter $\beta$ that captures the total probability of the channel gating in the system, defined as below:

$$\beta = \sum_{N=1}^{30} P_{opening}(N, \gamma) \cdot P_{formation}(N, \rho, \gamma) \tag{S12}$$

The probability of opening of a channel in a cluster of size $N$, $P_{opening}(N)$, is given as:

$$P_{opening}(N, \gamma) = a_1 \cdot (1 - a_2 \cdot e^{-C_1 \gamma}) e^{-C_2 N} \tag{S13}$$

The functional form was chosen to agree with the results of simulations given in Fig. S3, where a numerical fit gave $a_1 = 1.684$, $a_2 = 0.91$, $C_1 = 0.9274$, $C_2 = 0.49$. The *in vivo* values of membrane tension are calculated as before, from $\gamma = P \cdot r$ and equation S8. For a resting turgor pressure of 1 atm and cell radius of 0.5 um, the lateral tension on the cell membrane ranges from 50 to 80 mN/m for 15% volume increase during a hypoosmotic shock. Thus, the overall increase in membrane tension is 30 mN/m. Previous *in-vitro* report that the probability of MscL opening is half for a membrane tension ranging from 8-14 mN/m [9]. In order to scale the lateral membrane tension with the tensions in the coarse-grained model, we chose a mid-value of 11 mN/m.

Here it is important to keep in mind that our coarse-grained membrane model is phenomenological in nature, and while it produces the correct mechanical properties of biological membranes in general (bending rigidity and fluidity), it is not meant to reproduce the exact experimental membrane system. For instance, the membrane in our simulations ruptured at tensions greater than 3 mN/m, while experimentally reported values range between 1 mN/m and 25 mN/m, depending on the membrane composition and experimental conditions [10]. Moreover, our computer model does not include the presence of the bacterial wall, hence the value of the rupture tension in our generic model is not expected to match the experimentally reported values for bacterial envelope rupture. Nevertheless, the trends and physical mechanisms we observe will still be valid.

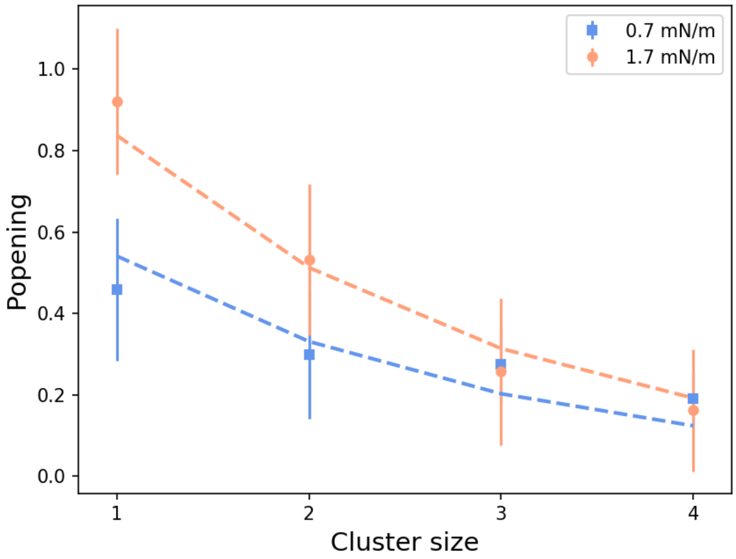


FIG. S3: **Probability of opening per channel versus cluster size at two values of membrane tension.** As the cluster size increases and the tension decreases, the probability of opening per channel decreases.

$P_{formation}(N, \rho, \gamma)$ is the probability of formation of cluster of size $N$ and at a given membrane tension and channel packing fraction, $\rho$, given as:

$$\rho = N_{MSC} \cdot a_{MSC} / A_c \tag{S14}$$

To obtain $P_{formation}(N, \rho, \gamma)$ we take into account that the computer model developed here, however minimal, does not allow us to collect good statistics for the cluster size distribution in a system of several hundreds of channels (and all at different tensions), which are the values reported in experiments [11]. Thus, to acquire better statistics for $P_{formation}$ we carry a separate set of simulations in which we model the channels as simple disks that interact with each other via a Lennard-Jones potential whose strength and range match the one measured in the membrane channel simulations, and at different values of membrane tensions. The disks were placed inside a simulation box and allowed to aggregate, similarly to how channels organise into clusters given enough time and a strong enough interaction ($\varepsilon_{protein-protein}$). This allowed an accurate estimation of the distribution of channels into clusters of different sizes. 1089 disks of diameter d $= 1\sigma$ were inserted inside a square box of side length $L$. The side length was varied from $L = 100\sigma$ to $L = 500\sigma$ in steps of $100\sigma$ and the 2D molecular dynamics simulations were carried out in the $NVT$ ensemble. The channels were allowed to interact through a Lennard-Jones potentials of depth $\varepsilon = 1.0kT$ to $\varepsilon = 2.0kT$ in steps of 0.2 kT. This range of interactions mimics the interactions observed within dimers of channels (Fig. S9). The disks were counted as being in the same cluster if found at distances $d \leq 3\sigma$, which corresponds to the distance at which the disks feel each others attraction in this simulation. The simulations were run for 5 million time steps each. The probability of formation for a cluster of size $N$ showed a non-linear dependence on the packing fractions (or cell volumes in the real system) as shown in Fig. S4. We can fit the observed probability of formation of a cluster of size N to a geometric distribution of the type:

$$P_{formation}(N, L, \varepsilon) = (1-p)^{N-1}p, \tag{S15}$$

where $p$ is the geometric distribution's ratio and related to the probability of formation of a monomer: $P_{formation}(1, L, \varepsilon) = p$. Equation S13 can be rewritten as follows:

$$lnP_{formation}(N, L, \varepsilon) = (N-1)ln(1-p) + lnp \tag{S16}$$

We assume that $p \propto L^2/\varepsilon$. The rationale for this is as follows: (i) if the system's size (for a fix packing fraction) increases, the probability of observing monomers will increase, and (ii) as the interaction energy between channels increases, they tend to reorganize into clusters and the probability of observing monomers will decrease. We can thus substitute $p$ for $p = mL^2/\varepsilon$, where $m$ is a parameter that captures the dependence of the geometric ratio, $p$, on $L^2$ and $\varepsilon$:

$$lnP_{formation}(N, L, \varepsilon) = (N-1)ln(1 - mL^2/\varepsilon) + ln(mL^2/\varepsilon) \tag{S17}$$

By using a Taylor approximation when $mL^2/\varepsilon << 1$:

$$lnP_{formation}(N, L, \varepsilon) = (N-1)(-mL^2/\varepsilon) + ln(mL^2/\varepsilon), \tag{S18}$$

which can be rewritten as

$$lnP_{formation}(N, L, \varepsilon) = (1-N)mL^2/\varepsilon + ln(L^2/\varepsilon) + lnm, \tag{S19}$$

Then, channel packing fraction, $\rho$, is inversely proportional to box area $L^2$:

$$\rho = \frac{N_{disks}\pi d^2}{4L^2}, \tag{S20}$$

So equation S17 becomes:

$$lnP_{formation}(N, \rho, \varepsilon) = (1-N)mN_{disks}\pi d^2/4\rho\varepsilon + ln(1/\rho\varepsilon) + ln(N_{disks}\pi d^2/4) + lnm \tag{S21}$$

Writing the equation S19 only in terms of the three variables $N$, $\rho$ and $\varepsilon$, it becomes:

$$ln[P_{formation}(N, \rho, \varepsilon)] = (1-N)A/\rho\varepsilon + ln(1/\rho\varepsilon) + B \tag{S22}$$

with $A$ and $B$ being constants to be estimated from the simulations. Fitting the results for $P_{formation}$ for both different $\rho$ and different $\varepsilon$ varied in the parameter space described above yielded the coefficients: $A = 0.009kT$ and $B = -8.316$.

We assume there is a correlation between the effective interaction energy between two channels with the surface tension, because higher surface tensions should lead to conformations that are more tilted and less likely to interact with each other. This is indeed observed in Fig. S9 which shows the average interaction energy between two channels if they are close enough from each other (explained in section II F). As the tension is increased, it is observed indeed that the channels tend to interact less strongly. We can then approximate the dependence of the interaction between channels on surface tension on a sigmoid (Fig. S9):

$$\varepsilon = a/(1 + \exp(-c(\gamma - d)) + b, \tag{S23}$$

where $a, b, c$ and $d$ are the parameters determined from the fitting the data in Fig. S9 ($a = 0.587kT$, $b = -1.973kT$, $c = 18.431$ m/mN, $d = 0.743$ mN/m).

We then normalise $P_{formation}$ as follows:

$$\sum_{N=1}^{30} P_{formation}(N, \rho, \gamma) = 1 \tag{S24}$$

We group together clusters larger than $N$=*30* as $P_{opening}$ and $P_{formation}$ are negligibly small for $N$>*30*, see Fig S3 and S4.

Lastly, based on our previous experimental results and cell volume recovery model, we scale the parameters of the coarse grained model in equation S22 and S23, such that at 4% cell volume increase all the channels are monomers [6] and a, b, c and d become $a = 19.371kT$, $b = -19.73kT$, $c = 1.675$ m/mN, $d = 3.715$ mN/m, respectively (Fig. S4).

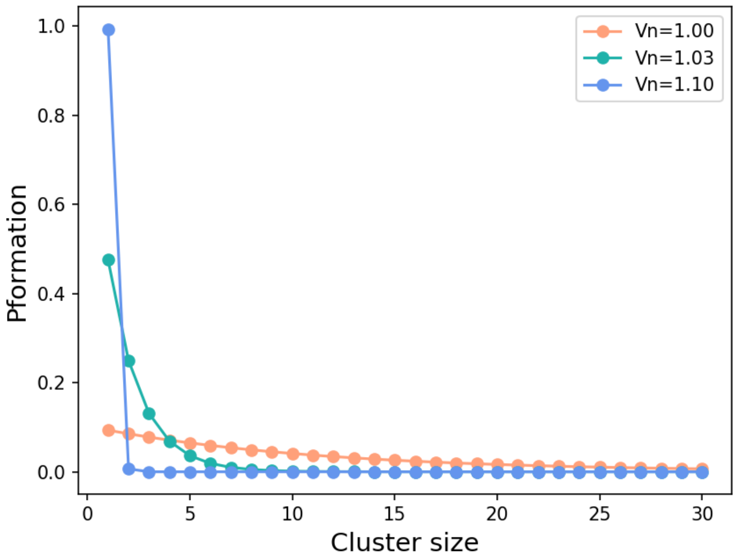


FIG. S4: **Probability of formation of a channel cluster at varying membrane tension.** The probability of formation of a cluster, calculated using Eq S22 at varying normalized cell volumes ($V_n$), at 0.5% packing fraction.

The Master equations for the cell volume change and solute concentration change (Eq. S10 and S11) can now be written as:

$$\frac{dV}{dt} = (\beta + 1)V_m KRT \left[ \left(\frac{n_i}{V} - c_e\right) - e^{\sqrt[3]{\frac{10}{3}}\pi \frac{E_0 l (V^{\frac{1}{3}} - V_0^{\frac{1}{3}})}{\Delta c_0 RTV^{\frac{1}{3}} V_0^{\frac{1}{3}}}} \cdot \Delta c_0 \frac{V_0^{\frac{1}{3}}}{V^{\frac{1}{3}}} \right] \tag{S25}$$

$$\frac{dn_i}{dt} = -\beta \cdot (V_m K \cdot \frac{n_i}{V} \cdot e^{\sqrt[3]{\frac{10}{3}}\pi \frac{E_0 l (V^{\frac{1}{3}} - V_0^{\frac{1}{3}})}{\Delta c_0 RTV^{\frac{1}{3}} V_0^{\frac{1}{3}}}} \cdot \Delta c_0 RT \frac{V_0^{\frac{1}{3}}}{V^{\frac{1}{3}}} + \alpha(\frac{n_i}{V} - c_0)). \tag{S26}$$

In Eq S26, $\alpha = D_s N_{MSC} a_{MSC} / l_M V_0$ . We fit the parameter values K and $\alpha$ to an experimental representative singe cell volume response as these two parameters are associated with the properties of channels and expect it to be different from the previous estimations [6] and fix the rest of the parameters at values previously determined (see section SI III for details and Supplementary Information of our previous model [6]). The 0.96 Osmol hypoosmotic shock was achieved by growing *E.coli* in media with elevated NaCl concentration [6] and then suddenly exposing the cells to the same media but without NaCl. Growing in elevated NaCl (specifically 550mM) results in expression of $\approx$ 1300 MscL [11]. The packing fraction at this experimental condition is 0.5%, calculated based on the area of a single MscL ($24nm^2$ [12]) and surface area of *E.coli* (6 $\mu$m$^2$ based on experimentally obtained cell volume of 1.338 fL [6]). The obtained fit values for K and $\alpha$ are $7.0 \times 10^{-22} molPa^{-1}s^{-1}$ and 0.37 $fls^{-1}$. To predict the cell responses when the channels existed solely as isolated channels or dispersed, $P_{formation}$ and $P_{opening}$ in Eq.S12 was calculated for $N = 1$.

#### IV. SUPPORTING RESULTS

##### A. Membrane tension

The calculation of the membrane tension revealed a linear scaling against the number of solute particles in the simulation box (Fig. S5). The membrane tension was fitted to a linear equation of the form $\gamma = aN_{solutes} + b$, with $a = 0.002$ mN/m and $b = -0.304$ mN/m. The membrane ruptured at tensions greater than 3 mN/m, corresponding to a number of solutes particles greater than 1800. This mapping of the surface tension was used in all subsequent calculations.

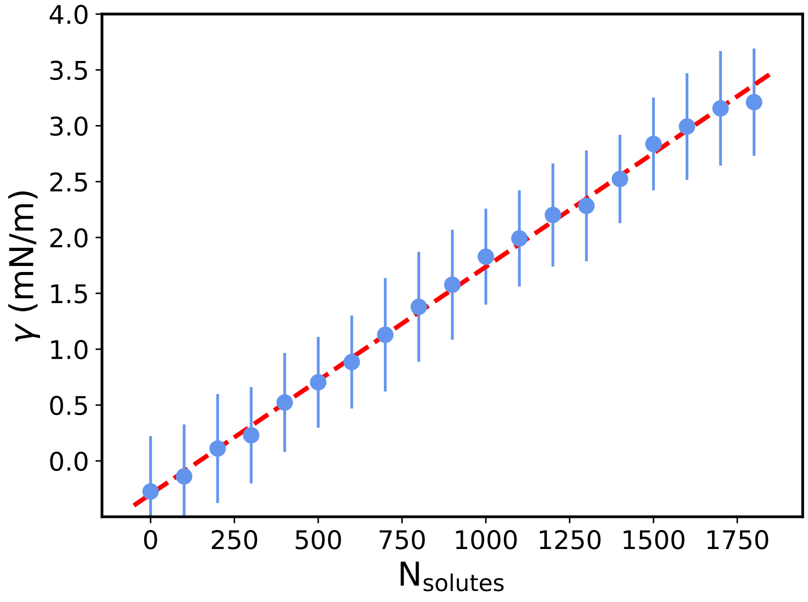


FIG. S5: **Membrane tension against number of solute particles.** The membrane tension scales linearly with the number of solute beads. The surface tension was calculated as an average over one million time steps. The error bars represent one standard deviation. Higher number of solutes (not shown here) led to the fracturing of the membrane.

##### B. Probability of opening

The opening probability of a single isolated MSC increases monotonically with membrane tension ( Fig. S5). This suggests the channel acts as a pressure sensor, more likely to be found in the "open" state at large external pressures. The threshold value of the gyration radius tensor term, above which the channel was considered as "open" in Fig. S6, was 8.2 nm$^2$.

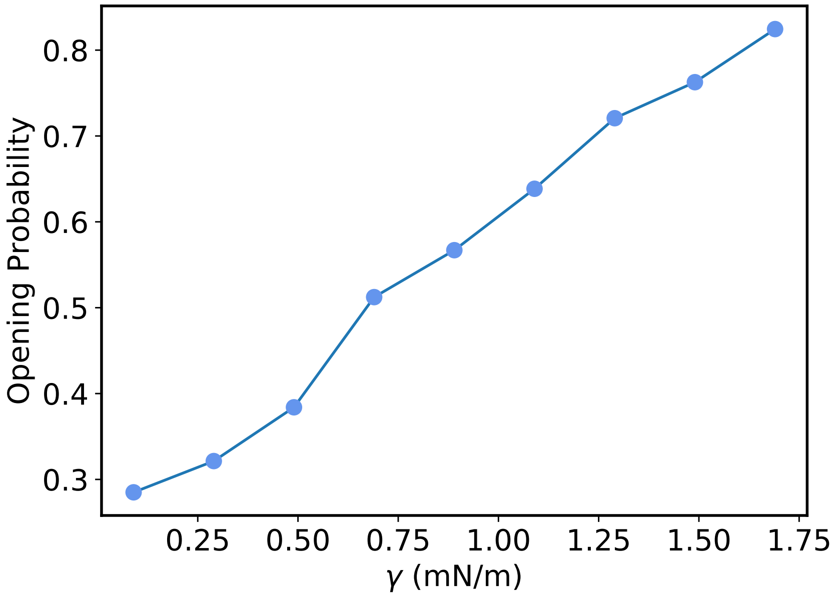


FIG. S6: **Opening probability of a single MSC against membrane tension.** The opening probability of a single isolated channel increases proportionally with the membrane tension. It should be noted that in the context of this model, there is still a finite probability for the channels to open even at low membrane tension. At higher tensions, the channels are more permeable to the solute particles, being found mostly in the open conformation.

##### C. Interaction of two channels

The potential of mean force (PMF) between two channels was calculated using the Weighted Histogram Analysis Method (WHAM) [13]. A biasing potential of a spring constant $k_{spring} = 1kT/\sigma^2$ was applied to the two channels and 20 different windows were used to extract the potential. In Fig. S7 it can be seen that two channels do not interact through any membrane mediated interactions, their only interaction being the volume exclusion at low separations.

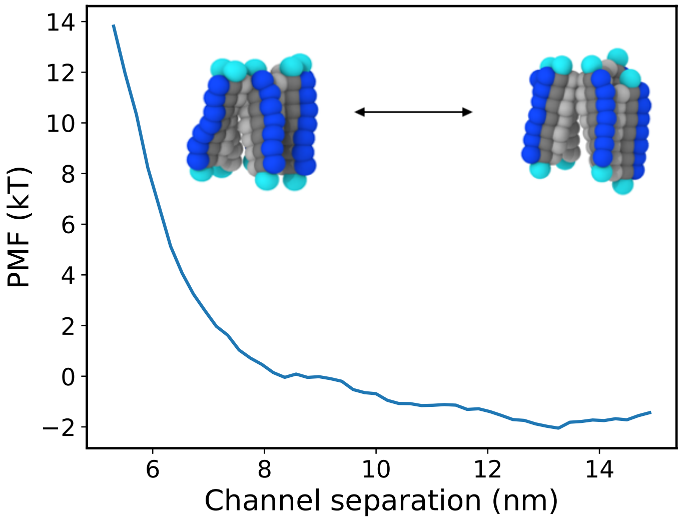


FIG. S7: **Potential of mean force between two non-interacting channels.** The potential of mean force between channels shows a relatively flat profile at large separations. The sudden increase at small separations is caused by the volume exclusion of the two channels.

##### D. Interaction of two rigid inclusions

Rigid inclusions of length 8 nm were inserted in the membrane to test the attractive effect of the hydrophobic mismatch. These inclusions were constructed similarly to the channels with a hydrophobic core to and head beads. The same potentials were used as for the MscL channels. The inclusions do not have any direct interactions. The potential of mean force (PMF) between two rigid inclusions was calculated from computing the radial distribution function of 10 channels placed on a lipid bilayer of side length $L_x = 40\sigma$, using the formula $PMF(r) = -k_B T ln(g(r))$. The inclusions did not organise into clusters, but showed a slight interaction of approximately 0.5kT (Fig. S8). It can be concluded that, although membrane-mediated interactions might be present in the system, they are too weak to drive the aggregation of channels and thus an attractive Lennard-Jones potential between channels was introduced to model the homophilic protein interactions. The simulations were run for 2 million time steps and 10 different seeds were used to compute the potential of mean force.

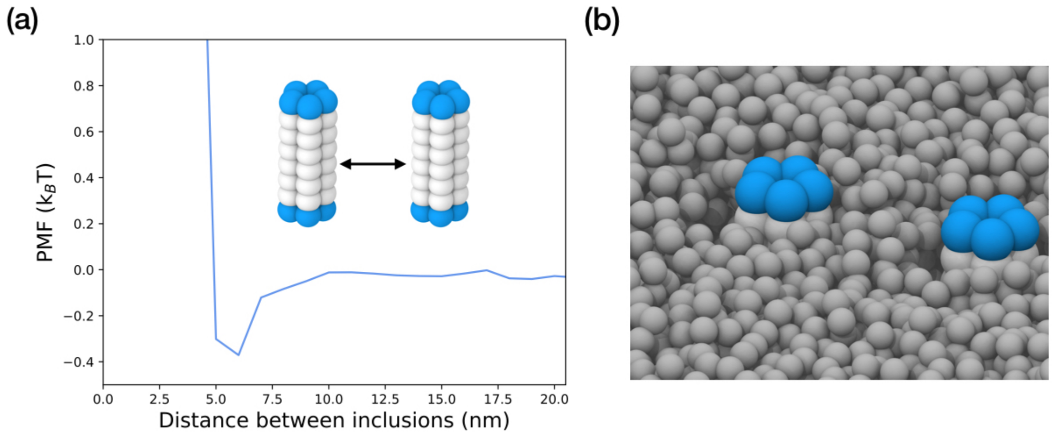


FIG. S8: **Potential of mean force between two rigid cylindrical inclusions of hydrophobic mismatch of 0.5 nm.** (a) The potential of mean force between two rigid cylindrical inclusions of a 0.5 nm hydrophobic mismatch shows weak attractive interactions of $\sim$ 0.5 kT. (b) Simulation snapshot of rigid inclusions embedded in the membrane.

##### E. Interaction of two attractive proteins

The probability of opening per channel is heavily influenced by the interaction of two adjacent channels. The increase in the direct protein-protein interaction leads to a decrease in the opening probability (Fig. S9). Higher interaction energies act to keep the two channels strongly bound together and allow for less conformational change. The channel has to overcome a larger activation energy barrier in order to rearrange from a compact closed structure into an expanded open one.

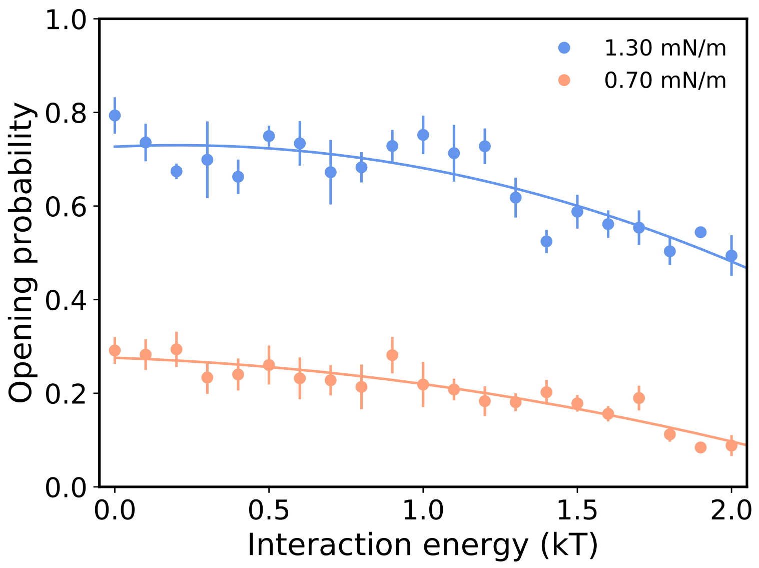


FIG. S9: **Opening probability for a channel against protein interaction energy $\varepsilon_{protein,protein}$.** The opening probability per channel of two interacting channels kept at a fixed distance from each other.

The interaction energy between two channels in a dimer was plotted in Fig. S10. The average interaction between channels decreases as the tension is increased. There is a sharp transition in the average interaction within a dimer at a tension of 0.7 mN/m. The behaviour can be attributed to tilting of the channels in the membrane plane at higher tensions, thus reducing the available contact area. Higher tensions will also suppress membrane thermal fluctuations, which induce the unbinding of two channels in a dimer.

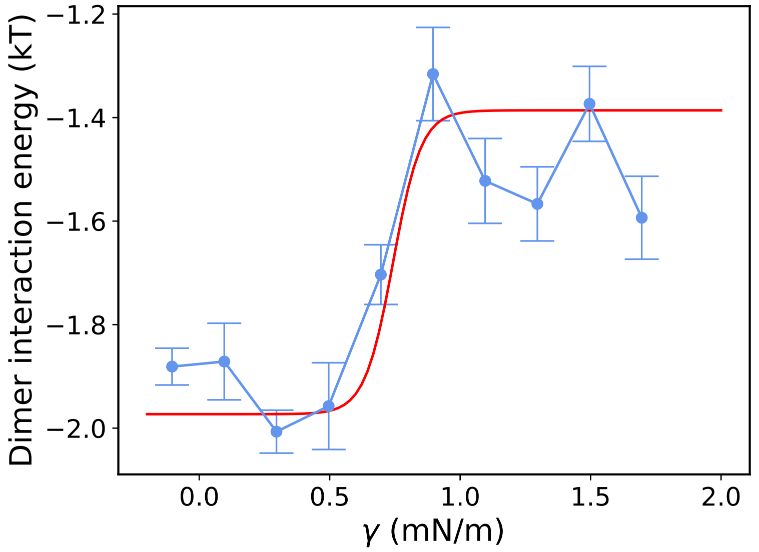


FIG. S10: **The average interaction energy between channels in a dimer depends on the membrane tension.** As the tension is increased, the channels tend to interact less favourably which leads to the disaggregation of protein clusters.

##### F. Model fit to experimental volume trace

We fit the master equations S25 and S26 to a representative experimental volume trace for 0.96 osmol [6] and physiological packing fraction of 0.5%, at this experimental conditions. The fitted trace was used to demonstrate the cluster aggregation-disaggregation dynamics as in the main text,Fig. 4a.

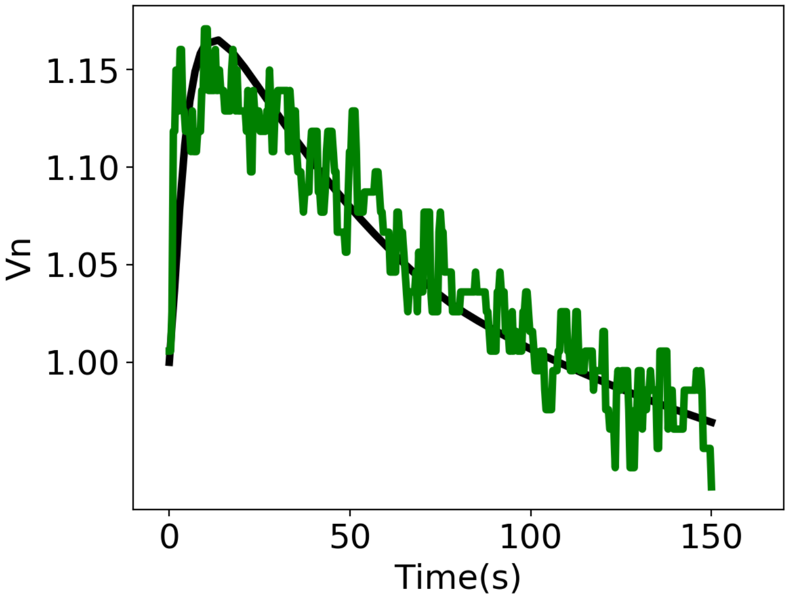


FIG. S11: Model fit (black) to a representative experimental volume trace (green) at 0.96 Osmol hypoosmotic shock and $\rho$=0.5%.

##### G. Dynamics of $\beta$ during hypoosmotic shock

The figure below demonstrates the dynamics of the effective probability of opening, $\beta$, (see Eq. S12) upon hypoosmotic shock, when the channels in the model are clustered and dispersed. Effective probability of opening $\beta$ in the clustered model shows an earlier drop in channel opening attributing to re-aggregation of channel clusters during volume recovery. This early closing of channels results in lesser volume overshoot while compared to the dispersed model (Fig4b).

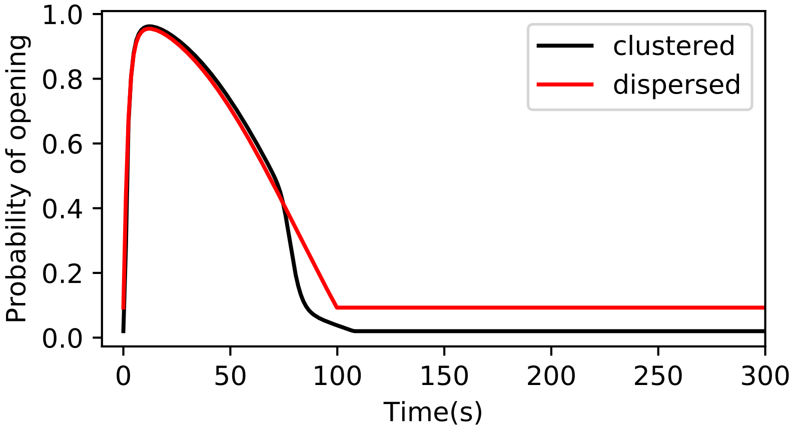


FIG. S12: Comparison of dynamics of the effective probability of opening, $\beta$, between a clustered model (black line) and dispersed model(red line) during a 0.96 Osmol hypoosmotic shock and 0.5% channel packing

##### H. Analysis of trade-offs different amounts of clustering pose on cell pressure and volume regulation

To illustrate the effects of increasing amount of clusters (higher than that predicted by coarse-grained model) on cell volume response (Fig 4b), interaction energy between the channels in Eq. S7 is increased by increasing the scaled constants a and b up to 20 times. The figure below shows the maximum tension on the cell membrane at maximum volume expansion, plotted against the minimum cell volume ($\Delta V_{n,min}$) during volume recovery. At a lower amount of channel clustering, the cells benefit by reducing the volume overshoot without much increasing the maximum tension upon a hyperosmotic shock. At higher amount of clustering, the benefit of further reducing $\Delta V_{n,min}$ is traded off by the burden on cells due to increased cell membrane tension, risking cell lysis. This illustrates that the interchannel interactions have been likely optimized to give the optimal amount of clustering.

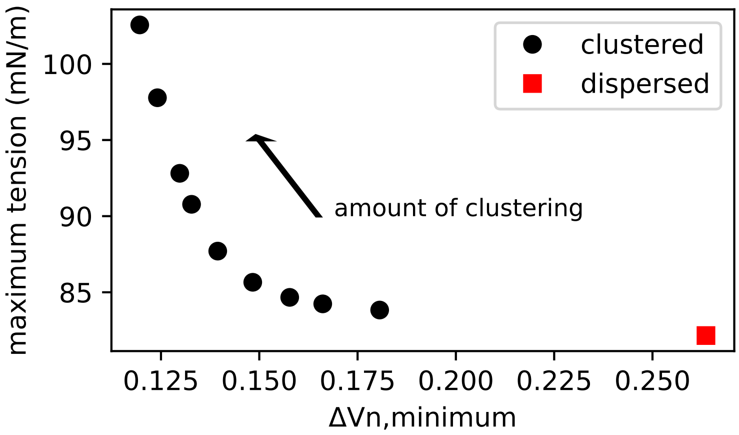


FIG. S13: **Effect of increased amount of clustering on the cell tension and overshoot upon a hypoosmotic shock.** Maximum tension on the cell membrane at maximum volume expansion is plotted against difference in volume overshot below the initial volume ($\Delta V_{n,min}$). The osmotic shock is 0.96 Osmol and packing fraction is set to 0.5%.

##### I. Supporting Videos

**Video 1:** Pore size oscillations of a single channel under osmotic shock leading to the membrane tension of 1.1 mN/m (see also Fig. 2b). Solute particles are omitted for clarity.

**Video 2:** Passage of a solute particle though a channel pore for membrane tension of 1.1 mN/m.

**Video 3:** Inter-protein attraction leads to channel clustering and cooperative closure; $\varepsilon_{protein,protein} = 0.9kT$ and $\gamma = 1.3$ mN/m (see also Fig. 3a). Solute particles are omitted for clarity.

**Video 4:** Side view: A preformed cluster made of 12 channels disaggregates under tension of $\gamma = 1.7$ mN/m (see Fig 3c). Here solute particles are explicitly shown (colored in pink).

**Video 5:** Top view: A preformed cluster made of 12 channels disaggregates under tension of $\gamma = 1.7$ mN/m (see Fig 3c). Solute particles are omitted for clarity.

[7] Yi Deng, Mingzhai Sun, and Joshua W. Shaevitz. Direct measurement of cell wall stress-stiffening and turgor pressure in live bacterial cells. 2011.
